# Metabolic and behavioral alterations associated with viral vector-mediated toxicity in the paraventricular hypothalamic nucleus

**DOI:** 10.1101/2023.10.26.564009

**Authors:** Rohan Savani, Erin Park, Nidhi Busannagari, Yi Lu, Hyokjoon Kwon, Le Wang, Zhiping P. Pang

## Abstract

**Objective:** Combining adeno-associated virus (AAV)-mediated expression of Cre recombinase with genetically modified floxed animals is a powerful approach for assaying the functional role of genes in regulating behavior and metabolism. Extensive research in diverse cell types and tissues using AAV-Cre has shown it can save time and avoid developmental compensation as compared to using Cre driver mouse line crossings. We initially sought to study the impact of ablation of corticotropin-releasing hormone (CRH) in the paraventricular hypothalamic nucleus (PVN) using intracranial AAV-Cre injection in adult animals.

**Methods:** In this study, we stereotactically injected AAV8-hSyn-Cre or a control AAV8-hSyn-GFP both Crh-floxed and wild-type mouse PVN to assess behavioral and metabolic impacts. We then used immunohistochemical markers to systematically evaluate the density of hypothalamic peptidergic neurons and glial cells.

**Results:** We found that delivery of one specific preparation of AAV8-hSyn-Cre in the PVN led to the development of obesity, hyperphagia, and anxiety-like behaviors. This effect occurred independent of sex and in both floxed and wild-type mice. We subsequently found that AAV8-hSyn-Cre led to neuronal cell death and gliosis at the site of viral vector injections. These behavioral and metabolic deficits were dependent on injection into the PVN. An alternatively sourced AAV-Cre did not reproduce the same results.

**Conclusions:** Our findings reveal that delivery of a specific batch of AAV-Cre could lead to cellular toxicity and lesions in the PVN that cause robust metabolic and behavioral impacts. These alterations can complicate the interpretation of Cre-mediated gene knockout and highlight the need for rigorous controls.

## Introduction

Located in the ventral forebrain, the paraventricular nucleus of the hypothalamus (PVN) acts as a major integrative center for diverse regulatory functions, including neuroendocrine response, autonomic tone, and complex behaviors including food intake and social interaction [1, 2, 3]. Dissecting the molecular, morphological, and functional heterogeneity of the PVN has thus far given rise to several distinct cell types involved in the control of these processes.

Refining our understanding of how the PVN integrates and outputs multimodal information across neuroendocrine and autonomic pathways may inform several disorders, including cardiovascular, metabolic, and neuropsychiatric disorders [2, 4, 5]. However, the PVN contains spatially intermingled but functionally distinct populations that can only be selectively targeted through genetic access, such as conditional knockout (KO) by Cre-lox recombination [6, 7]. While much work relies on promoter-driven Cre expression, another standard method relies on adeno-associated virus-mediated Cre expression (AAV-Cre) [8, 9, 10, 11]. Methods such as AAV-mediated Cre delivery in adult mice are valuable approaches to bypass transient developmental transcription and ensure high-fidelity expression in specific brain regions [8, 12]. However, emerging evidence suggests that there may be deleterious effects associated with AAV-Cre, resulting in morphological and behavioral abnormalities even in the absence of loxP sequences [13, 14].

In this study, we assessed the striking phenotype that emerged from the delivery of an AAV-Cre-prepared virus in the PVN of adult homozygous floxed corticotropin-releasing hormone (*Crh^f/f^*) mice. PVN CRH neurons are tightly linked to energy homeostasis—they respond to food cues, and acute manipulations decrease motivation to seek and consume food [8, 15, 16]. Chronic manipulations have been shown to confer risk to diet-induced weight gain, as well [17]. While knockout of CRH has not been shown to contribute to changes in metabolism or food intake thus far, these methods have used Cre driver lines or homologous recombination in embryonic stem cells [18, 19, 20]. As has been suggested for discrepancies in findings using knockout models of the type-1 CRH receptor, we hypothesized that developmental compensation may mask the metabolic effects of CRH knockout [21, 22]. We found that *Crh^f/f^* mice with putatively knocked out corticotropin-releasing hormone (CRH) via AAV-mediated expression of Cre recombinase in the PVN rapidly developed obesity and metabolic syndrome, which was an intriguing and unexpected finding. However, subsequent analyses for validation revealed that both CRH- and oxytocin-expressing neuron densities significantly decreased, likely due to cellular toxicity at the injection sites. Our study confirms the importance of the PVN in the control of metabolism and highlights the potential for toxicity due to a batch of a commonly used AAV, which necessitates careful interpretation and proper controls for experiments utilizing viral vectors.

### Hypothesis

We hypothesize that the targeted disruption of corticotropin-releasing hormone (CRH) expression within the paraventricular hypothalamic nucleus (PVN) through intracranial AAV8-hSyn-Cre injection in adult animals may result in obesity and metabolic alterations.

## Methods

### Animals

All experiments involving animals were conducted at the Child Institute of New Jersey, Rutgers University, USA. All protocols involving mice were approved by the Rutgers University Institutional Animal Care and Use Committee (IACUC) (Protocol # PROTO201702609) and by the National Institute of Health (NIH) guidelines. The animals used in this study were 5-7 weeks old at the time of stereotactic surgery. Homozygous *Crh^f/f^* mice (gift from Dr. Larry Zweifel) [23], *Glp1r^f/f^* mice [24], and wild-type (C57BL/6) mice were used in this study. For surgical injections, mice were anesthetized with isoflurane. For histological studies, mice were anesthetized with isoflurane and then followed by transcardial perfusion. For each experimental paradigm, littermate mice were randomized to each experimental group on the basis of body weight. Both males and females were used in this study. Mice of both groups were housed together. Investigators were blinded to treatment. The sample size required (n=5-7) for each group was estimated from previous studies [18].

### AAV vectors and stereotactic surgery

The AAVs used in this study included pAAV-hSyn-EGFP (Addgene, #50465-AAV8, titer: titer 7×10¹² vg/mL), pENN.AAV.hSyn.HI. eGFP-Cre.WPRE.SV40 (Addgene, #105540-AAV8, lot: v107437, titer: 1×10^13^ vg/mL), and AAV-hSyn-GFP-Cre (UNC Viral Vector Core, AAV8, lot: AV5053E, titer: 2.5 x 10^12^ vg/mL). For stereotactic injections, mice were anesthetized with isoflurane and placed in a stereotactic frame. Burr holes were drilled above the injection sites. 150 nL of the viruses were bilaterally stereotactically injected in the PVN (AP: −0.7, ML: ±0.18, DV: −4.7) at a rate of 1 nL/s (eGFP for control group, Cre for experimental group). For experiments in the dorsolateral septum (dLS), 100 nL of viruses were bilaterally injected in the dLS (AP: +0.5, ML: ±0.45, DV: −2.7) at a rate of 1 nL/second. Mice recovered for at least 3 weeks prior to conducting behavioral experiments. Injection sites were confirmed by *post hoc* histological examinations.

### Metabolic phenotyping

All experiments were conducted during the light cycle unless otherwise stated. Investigators were blinded to conditions during experiments.

#### Plasma corticosterone measurements

Blood was collected either at the beginning of the ITT test or during transcardiac perfusion. At least 10 μL was collected into EDTA-coated tubes and then centrifuged at 8000rpm at 4°C for 5 minutes. Extracted plasma was stored at −80° C until further processing. Corticosterone measurements were obtained with an ELISA as per manufacturer’s instructions (Enzo Life Sciences, ADI-900-097).

#### Food intake

Mice were singly housed before the experiment with ad libitum access to normal chow (Purina Mouse Diet 5058) and water. Chow remaining in the cage after each day (12 hr. light/dark cycle) was measured for 5 days and averaged together.

#### EchoMRI/fat mass measurements

Whole body composition and fat mass was measured by EchoMRI (Echo Medical Systems, Houston, Texas).

#### Comprehensive Lab Animal Monitoring System (CLAMS) assays

*Crh^f/f^* mice were placed into metabolic phenotyping cages (Columbus Instruments, OH, USA). Data was recorded every 15 minutes. Data was collected for 3 days (both light and dark cycles) after 2 days of habituation to single housing within the cages.

#### Glucose tolerance test (GTT) and insulin tolerance test (ITT)

Mice were fasted overnight (GTT) or for 6 hours (ITT). To minimize stress, 0.5 grams of chow was provided. For GTT, mice were intraperitoneally (IP) administered 20% glucose. Blood glucose measurements were collected at the 0, 15-, 30-, 60-, and 120-minute time points. For ITT, mice were IP-administered insulin. Injection volumes were 0.1 mL/10 g body weight. Blood glucose measurements were collected at the 0-, 15-, 30-, and 60-minute time points.

### Behavioral assays

#### Open field test

Mice were placed in the middle of a custom-made 45 x 45 cm square open field arena for 10 minutes. Analysis was performed using DeepLabCut v2.2 [25]. We labeled 150-200 frames taken from 14 or 10 videos, as resolution changed between recording sessions (95% of frames were used for training, 5% for testing). We used a ResNet50-based network with default parameters for 225,000 or 450,000 training iterations (until loss plateaued). After analyzing open-field videos, we then calculated the time spent in the inner zone (inner 3/5 of the arena, ∼25×25cm) using a region of interest tool. Distance traveled was calculated with custom code. Investigators were blinded to the condition during analysis.

#### Light-dark box test

Mice were placed in the SmartCage system (AfaSci) light side of a commercial light-dark box apparatus consisting of one dark and one illuminated compartment connected by a door. Time spent in each zone, latency to enter the dark zone, and number of entries were measured for 10 minutes. Data were analyzed using Excel.

### Histology and immunohistochemistry (IHC) analysis

Mice used for IHC analysis of CRH and oxytocin (OXT) were intracerebroventricularly administered colchicine a day prior to perfusion. Mice were anesthetized with isoflurane and perfused with PBS (pH 7.4) and then 4% paraformaldehyde (PFA). Coronal brain slices were cryosectioned (50 µm). For IHC analyses, brain slices were first incubated in blocking buffer (4% BSA, 1% goat serum in PBS) for 1 hour, then in primary antibody overnight at 4°C. Slices were washed with blocking buffer 3×15 min prior to incubation with a secondary antibody at RT for 2 hours, followed by further washing with PBS. Sections were mounted with Fluoroshield with DAPI (Sigma-Aldrich, # F6057). Images were collected with a Zeiss LSM700 confocal microscope. A Z-stack captured the entire thickness of the section with fluorescence. NeuN-stained slices were imaged at a 5 µm-thickness interval for a total range of 25 µm. Imaging analysis was performed using NIH-ImageJ and with the Allen Brain Atlas as a reference. The number of OXT- and CRH-cells in the PVN was counted with the Cell Counter plugin. To quantify NeuN density, images were background subtracted (20-pixel radius) before converting to a binary image. Then, segmented particles were filtered by despeckling twice and watershed transforming, followed by final counting (minimum size of 7 µm^2^). GFAP and Iba1 fluorescence area was calculated by dividing the area of thresholded signal by total PVN area. The quantifications were performed by two independent investigators who were blinded to the condition during analysis.

### Antibodies

The primary antibodies used in this study were rabbit Anti-CRH (1:500, BMA Biomedicals, #T-4037, RRID: AB_2314240), mouse Anti-OXT (1:500 Sigma-Aldrich, #MABN844), rabbit Anti-NeuN (1:500, Millipore, #ABN78, RRID: AB_10807945), rabbit Anti-GFAP (1:1000, Dako, #Z0334, RRID: AB_10013382), and chicken Anti-Iba1 (1:500, Aves Labs, #IBA1-0020).

### Statistical analysis

Statistical analyses were performed using GraphPad Prism 9. Data were analyzed using Student’s two-tailed t-tests and two-way ANOVA followed by Bonferroni’s multiple comparisons tests. Data are presented as mean ± SEM. A p-value < 0.05 was considered statistically significant.

## Results

### Delivery of AAV-mediated Cre in the PVN rapidly induces behavioral changes and metabolic syndrome in male Crh^f/f^ mice

Our initial goal was to apprise the functional role of CRH by knocking it out in the PVN in adult animals. To this end, we bilaterally stereotactically injected AAV8-hSyn-eGFP-Cre in the PVN of 5-7-week-old *Crh^f/f^* mice, with AAV-hSyn-eGFP (Control) injected in littermate controls (**Fig. 1A**). After four weeks of recovery, we found that plasma corticosterone levels in Cre-injected *Crh^f/f^* mice were markedly reduced, which was highly suggestive of possible loss of function of CRH (**Fig. 1B**). Even with normal chow feeding, the Cre-injected mice showed robust and rapid body weight gain within days. Cre-injected mice reached an average of 50.5 ± 0.7 grams in 6 weeks, while the average body weight of control animals was 28.2 ± 0.7 grams (**Fig. 1C**). The increased body weight in the Cre-injected group might be, at least in part, due to elevated food intake (**Fig. 1D**). We repeated these experiments in an independent cohort of *Crh^f/f^* female mice (**Fig. S1A**). In this, the Cre-injected group fed with normal chow weighed 45.2 ± 4.2 grams 6 weeks after viral injection (**Fig. S1B**), indicating that the changes observed thus far occur in a sex-independent manner.

**Figure 1.**
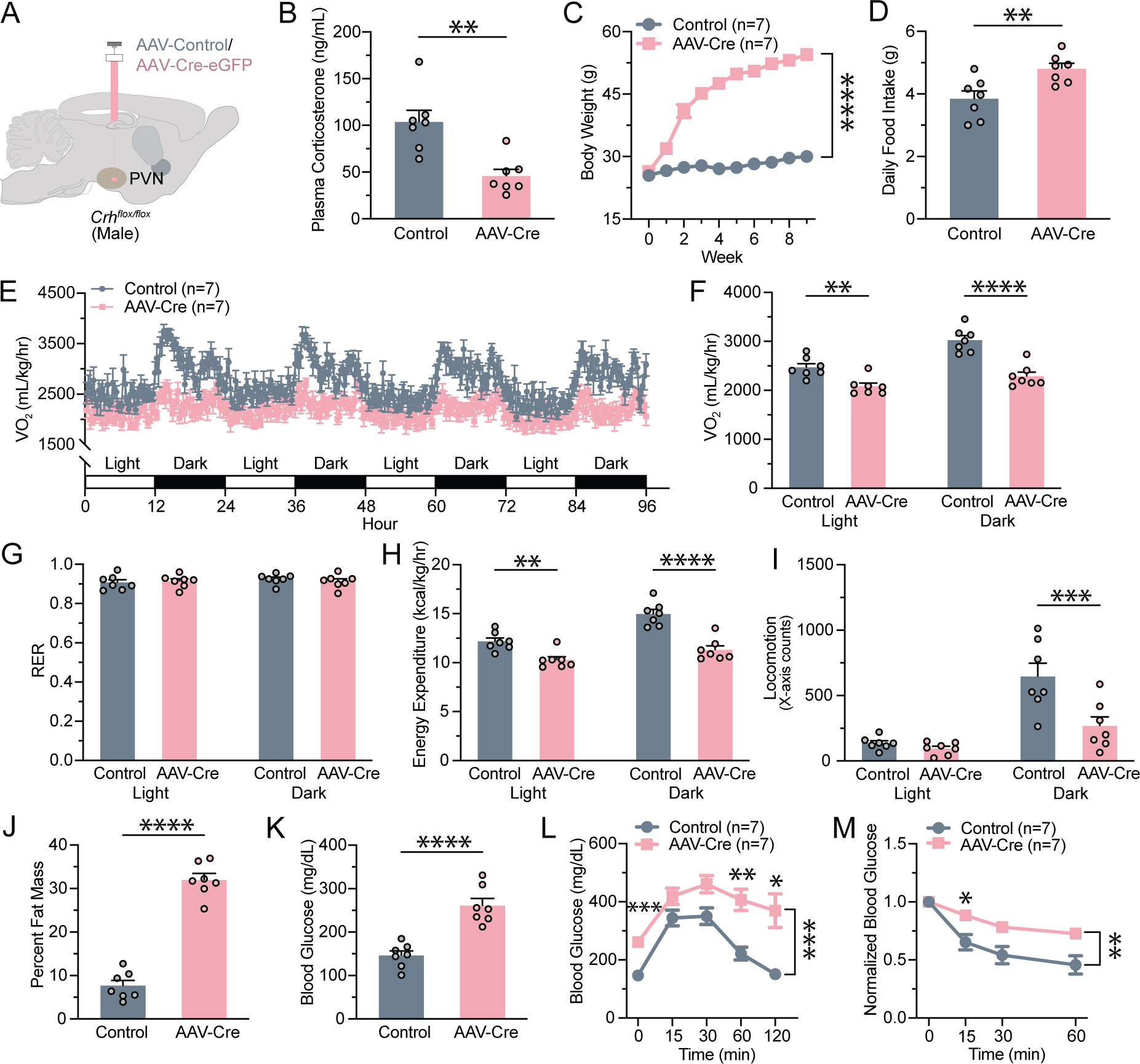
AAV-Cre administration in PVN of male *Crh^f/f^* mice results in the development of metabolic syndrome. **(A)** Experimental paradigm for virus delivery. AAV-Cre or AAV-eGFP was stereotactically injected into the adult male murine PVN. **(B)** Plasma corticosterone decreased following AAV-Cre injection (t (12) = 4.011, p = 0.0017). **(C)** Changes in body weight following virus delivery at week 0 (two-way ANOVA, main effect of Group: F (1, 12) = 294.1, p < 0.0001; main effect of Time: F (1.378, 16.53) = 228.5, p < 0.0001; interaction between Group and Time: F (9, 108) = 138.4, p < 0.0001). **(D)** Daily chow intake following virus injection increased with AAV-Cre (t (12) = 3.152, p = 0.0083). **(E)** Changes in oxygen consumption were observed over the course of 96 hours. **(F)** Quantification of the decrease in oxygen consumption in AAV-Cre-injected mice (two-way ANOVA with Bonferroni’s multiple comparisons test, main effect of Group: F (1, 12) = 25.11, p = 0.0003; main effect of Time: F (1, 12) = 208.4, p < 0.0001; interaction between Group and Time: F (1, 12) = 43.58, p < 0.001). **(G)** The respiratory exchange ratio (RER) was unaltered (two-way ANOVA, no main effect of Group: F (1, 12) = 0.02021, p = 0.8893; no main effect of Time: F (1, 12) = 2.970, p = 0.1105; no interaction between Group and Time: F (1, 12) = 3.566, p = 0.0834). **(H)** Energy expenditure decreased following AAV-Cre delivery (two-way ANOVA with Bonferroni’s multiple comparisons test, main effect of Group: F (1, 12) = 26.31, p = 0.0002; main effect of Time: F (1, 12) = 207.9, p < 0.0001; interaction between Group and Time: F (1, 12) = 45.10, p < 0.0001). **(I)** Locomotion in the x-axis in the CLAMS cage decreased during the dark cycle (two-way ANOVA with Bonferroni’s multiple comparisons test, main effect of Group: F (1, 12) = 8.461, p = 0.0131; main effect of Time: F (1, 12) = 41.86, p < 0.0001, interaction between Group and Time: F (1, 12) = 10.17, p = 0.0078). **(J)** Proportion of fat mass as determined by EchoMRI increased in AAV-Cre-injected mice (t (12) = 12.67, p < 0.0001). **(K)** Changes in average blood glucose after overnight fasting (t (12) = 5.888, p < 0.0001). **(L)** The glucose tolerance test reveals increased baseline blood glucose levels (two-way ANOVA with Bonferroni’s multiple comparisons test, main effect of Group: F (1, 12) = 22.30, p = 0.0005; main effect of Time: F (2.717, 32.60) = 24.72, p < 0.0001; interaction between Group and Time: F (4, 48) = 3.111, p = 0.0235). **(M)** The insulin tolerance test reveals insulin tolerance (two-way ANOVA with Bonferroni’s multiple comparisons test, main effect of Group: F (1, 11) = 11.30, p = 0.0063; main effect of Time: F (1.712, 18.84) = 55.72, p < 0.0001; no interaction between Group and Time: F (2, 22) = 0.6622, p = 0.5257). Sample sizes are as shown on graphs (n = 7 mice per group). Data are presented as individual points and mean ± or + SEM.

As PVN-derived CRH is known to be involved in stress-related behaviors, coping, and relief [16, 26, 27, 28], we hypothesized that reduced CRH release and corticosterone levels (**Fig. 1B**) in the Cre-injected animals would be anxiolytic. To test this hypothesis, we examined anxiety-like behavior in AAV-Cre injected *Crh^f/f^* male mice in the open-field and light-dark box assays (**Fig. S2**). Contrary to what we hypothesized, we found that the Cre-injected mice spent less time in the center zone in the open-field assay (**Fig. S2B**), and they spent more time in the dark zone during the light-dark box test (**Fig. S2D**). These tests suggested that AAV-Cre in the adult PVN may lead to heightened anxiety-like behaviors.

Nevertheless, the drastic increase in body weight of AAV-Cre-injected animals prompted us to further assess their metabolic phenotype. We used CLAMS to measure energy intake and expenditure, as well as to identify any changes in nutrient utilization. Compared to AAV-GFP-injected animals, AAV-Cre-injected mice consumed less oxygen throughout the entire day (**Fig. 1E-F**). The respiratory exchange ratio was equal between both groups, indicating no shift in nutrient utilization accompanying increases in weight (**Fig. 1G**). Energy expenditure was significantly decreased in AAV-Cre-injected mice in both light and dark cycles (**Fig. 1H**). This may be in part due to a decrease in locomotor activity (**Fig. 1I**). These data suggest that alterations in energy expenditure may contribute to the development of obesity alongside hyperphagia in AAV-Cre-injected mice. Consistent with these findings, Cre-injected mice display significantly increased body fat composition (**Fig. 1J**) and significantly increased fasting plasma glucose levels measured (**Fig. 1K**). To evaluate any changes in glucose and insulin homeostasis, we also performed GTT and ITT. GTT revealed that Cre-injected mice had consistently elevated glucose levels, indicative of higher glucose tolerance (**Fig. 1L**), while ITT revealed that they had insulin resistance (**Fig. 1M**). These results indicate that Cre recombinase expression in the PVN region of *Crh^f/f^* mice leads to obesity development and maintenance through changes in food intake behavior and energy expenditure and causes metabolic syndrome resembling type 2 diabetes mellitus.

Having observed the development of diabetes and obesity in the PVN Cre-injected *Crh^f/f^* mice, we sought to validate these findings as attributable to the depletion of hypothalamic CRH. We examined CRH expression using immunohistochemistry and found decreased CRH-positive cell density in the PVN of AAV-Cre-injected mice (**Fig. 2A & 2C**). This suggested that CRH expression was suppressed, which supports the hypothesis that PVN CRH participates in regulating metabolism. However, this hypothesis is complicated by the observation of decreased OXT expression in brain sections of the same animals with Cre injections (**Fig. 2B & 2D**). Decreased expression of both CRH and OXT was accompanied with a trend toward reduced cell density as measured by DAPI staining (**Fig. 2E**). Based on these data, we caution that virally expressed Cre recombinase may be sufficient to deplete CRH from the PVN of *Crh^f/f^* animals but may also have an off-target impact on PVN neuronal cells.

**Figure 2.**
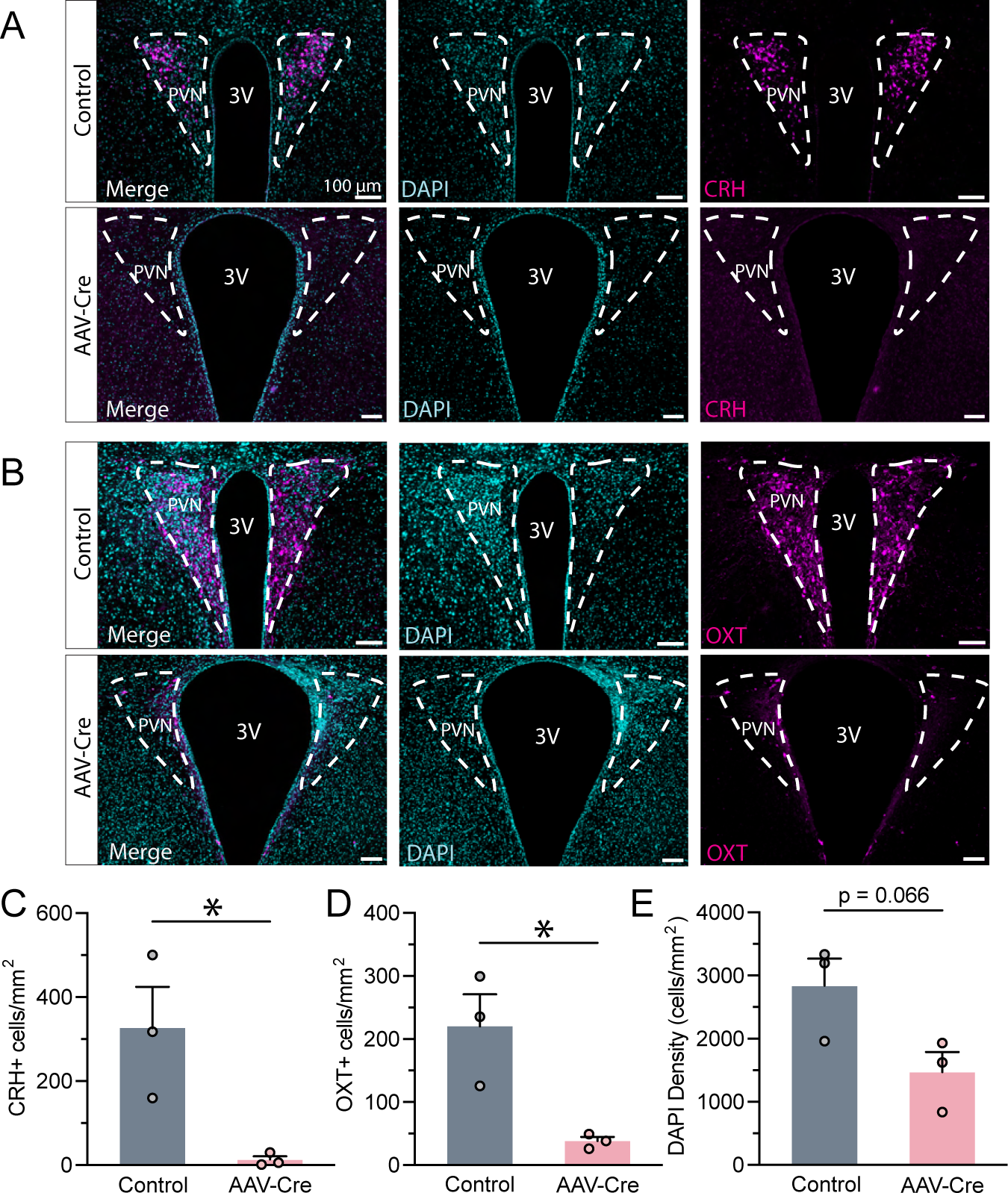
AAV-Cre action in PVN of *Crh^f/f^* mice depletes CRH and OXT. **(A)** Example comparison of CRH expression in PVN between both groups. **(B)** Example of OXT expression in PVN. **(C)** CRH-positive cell density decreased following AAV-Cre expression in *Crh^f/f^* PVN (t (4) = 3.176, p = 0.0337). **(D)** OXT-positive cell density decreased as well (t (4) = 2.807, p = 0.0236). **(E)** Overall cell density trended toward a decline (t (4) = 2.511, p = 0.0660). Sample sizes are as shown on graphs (n = 3 mice per group). Data are presented as individual points and mean + SEM.

### Delivery of AAV-mediated Cre in the PVN rapidly alters behavior and causes metabolic syndrome in wild-type animals

While OXT expression may have decreased due to the development of obesity, the trend of decreased cell density in the AAV-Cre group led us to hypothesize that the AAV-Cre virus may impact PVN neuronal survival. The toxicity of Cre expression on cellular health has been reported previously [13, 14, 29, 30].

To this end, we examined the impacts of AAV-Cre in C57BL/6 wild-type (WT) mice (**Fig. 3A**). Similar to the AAV-Cre injected *Crh^f/f^* animals, we found a marked reduction in plasma corticosterone levels in WT mice injected with AAV-Cre (**Fig. 3B**). They also gained significant body weight potentially due to hyperphagia (**Fig. 3C-D**). Given that our observations closely mirrored those found in *Crh^f/f^* mice, we also assessed changes in anxiety-like behaviors (**Fig. S3A & S3C**). AAV-Cre-injected mice spent less time in the center zone during the open field test (**Fig. S3B**). Consistent with this, Cre mice spent more time in the dark zone of the light-dark box (**Fig. S3D**). These data suggest that the behavioral, metabolic, and cellular alterations observed with Cre injection in the PVN may be due to the expression of the Cre-encoding viral vector.

**Figure 3:**
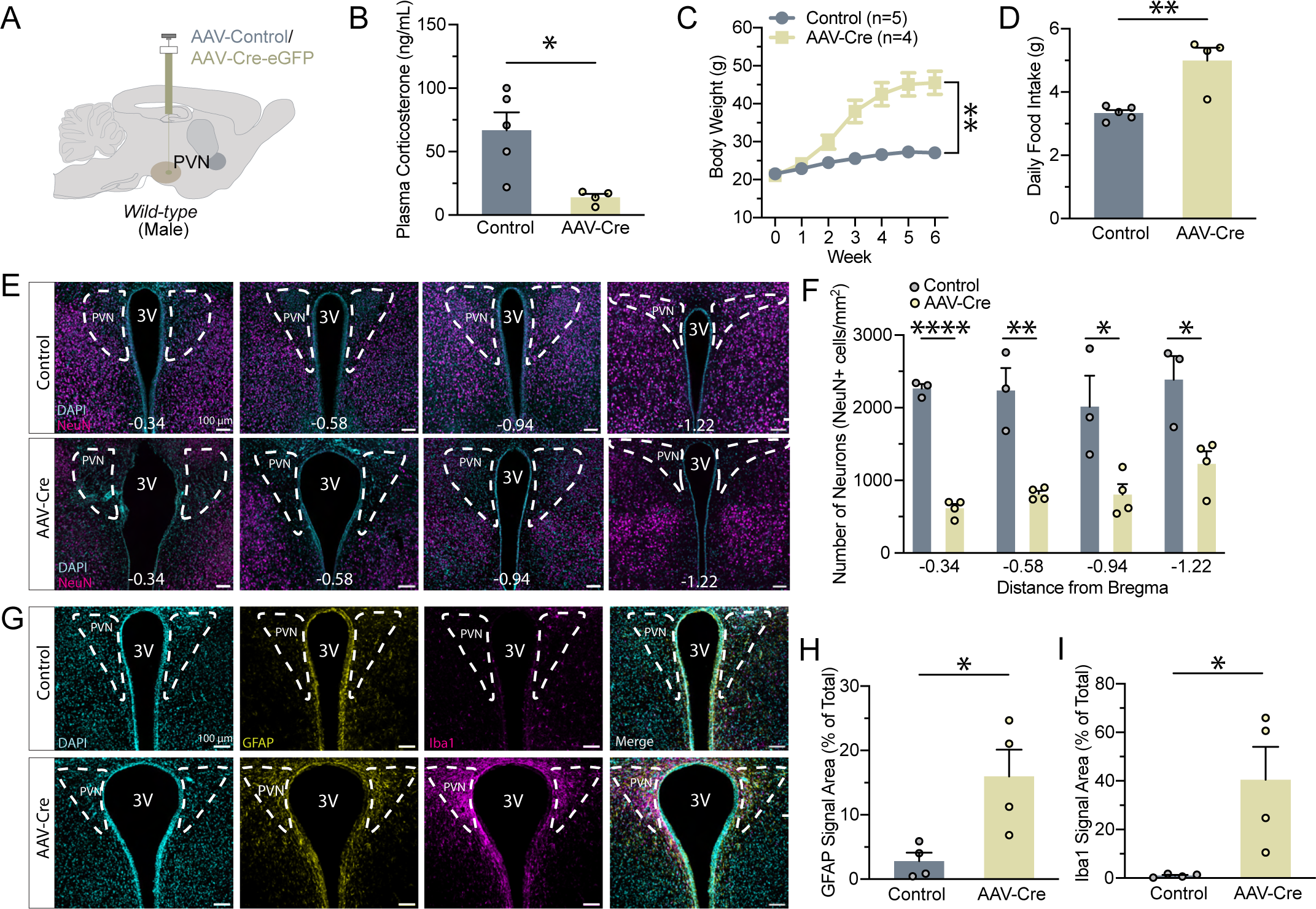
AAV-Cre action in wild-type PVN induces metabolic changes, neuronal cell death, and recruitment of glia. **(A)** Experimental paradigm for virus delivery in wild-type male mice. **(B)** Plasma corticosterone levels decreased after AAV-Cre injection (t (7) = 3.260, p = 0.0139). **(C)** Body weight changes followed virus delivery (two-way ANOVA, main effect of Group: F (1, 7) = 22.37, p = 0.0021; main effect of Time: F (1.134, 7.940) = 191.5, p < 0.0001; interaction between Group and Time: F (6, 42) = 79.71, p < 0.0001). **(D)** Daily food intake increased in AAV-Cre-injected mice (Mann-Whitney test, U = 0, p = 0.0159). **(E)** Example images of neuronal nuclei (NeuN) across the PVN following injection in wild-type mice. **(F)** Quantification of neuron density at different stereotactic coordinates: −0.34 (t (5) = 18.72, p < 0.0001), −0.58 (t (5) = 5.343, p = 0.0031), −0.94 (t (5) = 3.053, p = 0.0283), and −1.22 (t (5) = 3.372, p = 0.0199). **(G)** Example comparison of glial fibrillary acidic protein (GFAP) and ionized calcium binding adaptor molecule 1 (Iba1) expression in the wild-type PVN between the two groups. **(H)** Expression of astrocytes increased as quantified by GFAP fluorescence area (t (6) = 3.017, p = 0.0235) **(I)** Expression of microglia increased as quantified by Iba1 fluorescence area (t (6) = 2.917, p = 0.0267). Sample sizes are as shown on graph: **(F)** n = 3 control, 4 AAV-Cre **(H, I)** n = 4 per group. Data are presented as individual points and mean ± or + SEM.

We next systematically evaluated the number of neurons in the PVN of WT mice following AAV-Cre or AAV-GFP injections using neuronal nuclei (NeuN) staining (**Fig. 3E**). Expression of NeuN was significantly decreased across most of the PVN with the administration of AAV-Cre, likely due to diffusion (**Fig. 3F**). Again, accompanied by the enlarged visual appearance of the third ventricle, the widespread decline in neuronal density in AAV-Cre-expressing mice lends support to the idea of toxic and nonspecific AAV-Cre action, resembling the effects of a chemical lesion for PVN neuronal cells [31, 32]. This neuronal cell loss is accompanied by astrogliosis and microglial cell activation (**Fig. 3G-I**).

Together, these data suggest that AAV-Cre injection may cause non-specific cell death at the injection site, with PVN neuronal ablation causing obesity and diabetes. This is consistent with previous reports that PVN lesions result in severe obesity and dysregulated metabolism [31, 32, 33, 34, 35].

### Delivery of AAV-mediated Cre in the dorsal lateral septum induces neuronal cell lesions without causing metabolic syndrome in Glp1r ^f/f^ animals

To further confirm the apparent toxicity of the AAV-Cre that we used, we tested it in another brain region: the dorsal lateral septum (dLS). The glucagon-like peptide-1 receptor (GLP-1R) is highly expressed in dLS and is implicated in feeding and metabolism [36, 37, 38, 39]. We assessed if a diluted AAV-Cre could be used to determine endogenous GLP-1R function. To prevent the possibility of cellular toxicity due to high AAV titers, we diluted our AAV to 5 x 10^12^ vg/mL and only injected 100 nL. We injected this AAV-Cre or a control vector in *Glp1r ^f/f^* mouse dLS (**Fig. S4A**), but we did not observe significant changes in body weight or metabolism within 4 weeks of injection (**Fig. S4B**). However, we found significantly decreased NeuN positive cell density (**Fig. S4C-D**), suggesting that Cre viral vector injection in the mouse brain causes neuronal cell loss.

### Delivery of an alternative preparation of AAV-Cre in PVN avoids cell death and body weight changes in Crh^f/f^ mice

We next tested if our findings were representative of all viral vectors delivering Cre recombinase. We injected a preparation of AAV8-hSyn-Cre-GFP (UNC Vector Core) at the provided titer (2.5 x 10^12^ vg/mL) in Crh^f/f^ male mouse PVN (**Fig. 4A**). We found minimal changes in body weight after 6 weeks of injection (**Fig. 4B**), suggesting that toxicity was attenuated. Accordingly, we found no changes in neuronal cell density (**Fig. 4C-D, Fig. S5A&B**). Together, these results point toward disparate and variable effects of AAV-Cre viruses, potentially due to preparation or construction of the specific batch of viruses.

**Figure 4:**
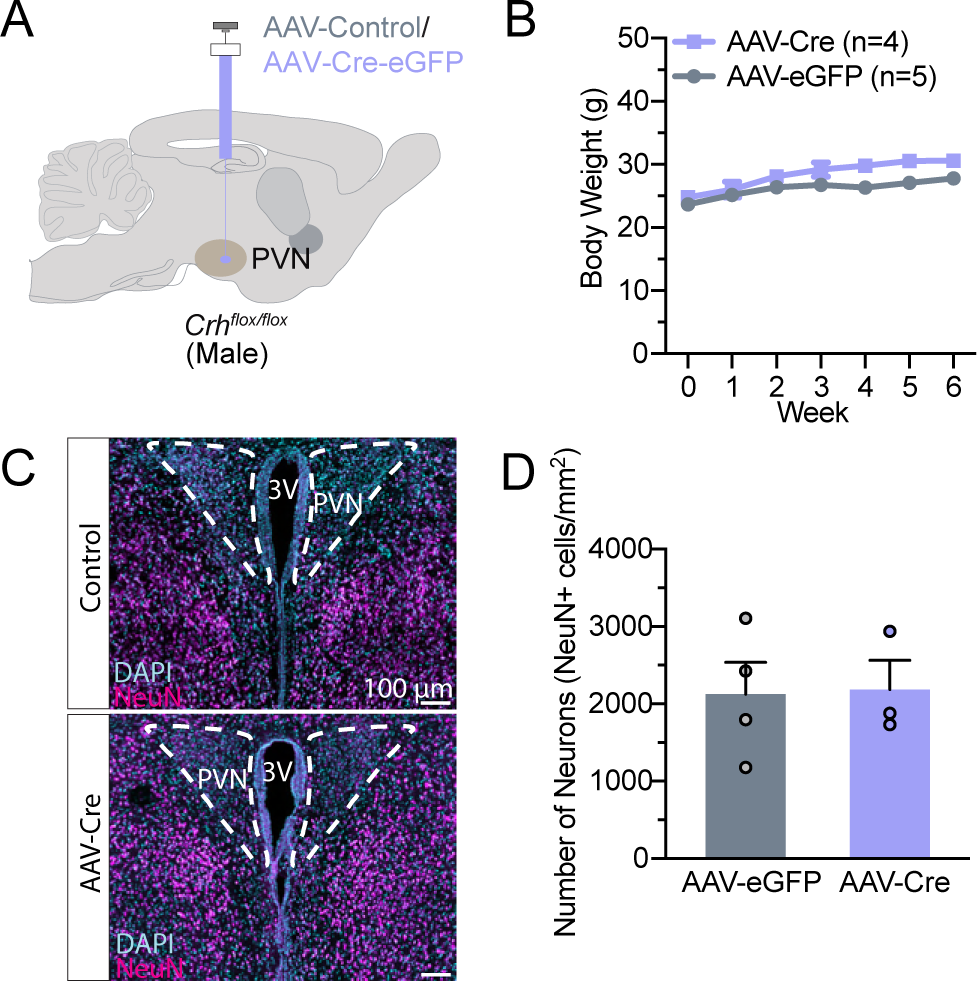
An alternative preparation of AAV-Cre does not lead to cell death in PVN. **(A)** Experimental paradigm for virus delivery in male *Crh-flox* mice. (B) Body weight does not change following delivery (two-way ANOVA, main effect of Group: F (1, 7) = 5.304, p = 0.0547; main effect of Time: F (6, 42) = 19.71, p < 0.0001; interaction between Group and Time: F (6, 42) = 1.729, p = 0.1379). **(C)** Example images of NeuN density in PVN following injection. **(D)** Neuron density did not change following AAV-Cre injection (t (5) = 0.1005, p = 0.9238). Data are presented as individual points and mean ± or + SEM.

## Discussion

In this study, we showed that delivery of one preparation of AAV-Cre recombinase in the PVN of adult mice can lead to the development of metabolic syndrome and alterations in food-related and anxiety-like behaviors, as well as metabolic phenotype. We demonstrated that *Crh^f/f^* mice expressing AAV-Cre in PVN develop obesity largely mediated by hyperphagia and changes in energy expenditure. Moreover, this develops alongside glucose tolerance and insulin resistance, as well as decreased locomotion. We investigated whether CRH was truly and specifically depleted and found off-target effects on OXT expression (**Fig. 2**) and cell loss at sites of Cre viral vector injections in either *Crh^f/f^* or WT mice. These data suggested off-target effects of our viral vector and indicate that caution should be used when interpreting the physiological consequences of AAV injection in adult animals. Intriguingly, we found that another preparation of AAV-Cre did not lead to lesions in PVN. Differences in how these viral vectors (batches, lots, or other procedural variations) were constructed or prepared may potentially be the source of this disparity. Variability in viral vector quality further accentuates the need to validate reagents prior to experimental use.

Specific depletion of PVN-derived CRH has thus far not been shown to induce massive metabolic dysregulation when knocked out using Cre expression driven by either *Crh^f/f^* crossed with *Sim1-Cre* mice or CRISPR [18, 40]. Our initial goal was to assess the impact of ablation of PVN CRH in adult animals by using stereotactic injection of AAV-Cre. We observed massive obesity and metabolic syndromes that starkly differ from other models of CRH signaling deficiencies [18, 20, 40, 41, 42, 43]. CRH-deficient mice have intact feeding behavior, even after stress [41]. Deletion of the type-1 CRH receptor yields conflicting results on food intake but largely minimal effects on body weight [22, 44]. Global deletion of the type-2 receptor similarly results in no changes in body weight despite significant increases in food consumption, likely due to increased energy expenditure and sympathetic tone [45]. Despite the observation of reduction of CRH expression and plasma corticosterone levels, which may be an indication of impaired regulation of the hypothalamic-pituitary-adrenal axis (HPA), we surmise that any metabolic or behavioral deficits reflected off-target effects of AAV-Cre. Indeed, we confirmed that neuronal cell death and reactive gliosis occurred in the PVN with AAV-Cre expression, resembling a response to central nervous system injury. Therefore, the study cannot conclude whether adult ablation of CRH in the PVN leads to metabolic or behavioral changes in adult animals. While we have preliminarily shown that an alternatively sourced AAV-Cre does not cause significant body weight changes in *Crh^f/f^* mice, further behavioral and metabolic phenotyping—as well as validation that CRH is fully knocked out—must be performed in future studies to concretely resolve this aim.

The PVN is speculated to provide top-down control of feeding-related information processing via intrahypothalamic connections or dense projections to midbrain and hindbrain regions [2]. These results could not provide insights on CRH impact on metabolism but highlight the necessity of the PVN in regulating energy homeostasis and body weight. These findings are in accord with prior ablation experiments and more recent work on genetic knockout models specific to the PVN [31, 32]. It has been shown that unilateral PVN electrolytic lesions would increase lipid accumulation in white adipose tissue ipsilateral to the side of the PVN [46]. Chronically inhibiting PVN MC4R/PDYN neuron synaptic release by expressing tetanus toxin or ablating those neurons by expressing caspase-3 resulted in pronounced hyperphagia, obesity, and a significant elevation in food consumption [32]. Our metabolic phenotyping data revealed that cellular lesions of PVN develop a strong obesity phenotype largely mediated by hyperphagia and changes in energy expenditure. Moreover, dysfunction of the PVN induces hypersomnia and mediates the diurnal rhythm of metabolism in mice [47, 48]. Consistent with this, we also found that changes in energy expenditure were more pronounced in AAV-Cre-injected mice during dark cycles (**Fig. 1H**).

Manipulating gene expression on and off using AAVs in genetically engineered animals is one of the most powerful tools to bypass transient developmental transcription for studying gene function *in vivo* [8, 12]. For example, it has been shown that knockout of *Glp1r* via AAV-Cre in PVN demonstrated a significant obesity phenotype, which was not observed in *Glp1r^f/f^* mice cross with SIM1- or NKX2.1-Cre mice [8, 49]. Similarly, AAV-CRISPR–Cas9-mediated ablation of leptin receptor in arcuate nucleus demonstrated that agouti-related peptide (AgRP) neurons but not proopiomelanocortin (POMC) neurons are required for the primary action of leptin to regulate both energy balance and glucose homeostasis [12]. The present results add to a growing body of literature suggesting that toxicity occurs irrespective of serotype and brain region, though a systematic comparison is lacking. While we show toxicity with one viral preparation in this study, previous studies have reported varying levels of success with this AAV-Cre and others [9, 13, 14, 48, 50]. In the mesolimbic reward system, it has been reported that Cre expression in the ventral tegmental area and substantia nigra can induce toxicity associated with perturbations of dopamine-related behaviors [13, 14]. Further work delineating different viral preparations and optimal titers for neuronal manipulations will be needed. Inducible systems of Cre expression, such as those using tamoxifen, offer an appealing alternative to constitutively expressed Cre systems; however, administration of tamoxifen has also been reported to lead to cytotoxicity and off-target effects [51].

Our data confirm that the PVN is an important brain region in the regulation of metabolism; dysfunction of the PVN, such as via neuronal death, causes obesity and type 2 diabetes. We also conclude that AAV-Cre vector toxicity ablates PVN neurons and results in a phenotype that is not associated with the gene of interest. These results underscore the importance of rigorous controls to ensure virus quality and fidelity of action. Caution must especially be exercised when the observed phenotype closely matches that of lesions in the area of use. Ensuring that expression in wild-type mice results in no observable changes or that neuronal cell densities are unaltered (if they are not expected to be) are two such methods of validating that the AAV is not toxic on its own. Together, viral vectors represent a class of worthwhile tools to manipulate the nervous system; however, scientists must take care to ensure that the potential for toxicity in commonly used AAVs is ruled out in experimental design and data interpretation.

## Future Directions

In this study, we exclusively examined the impact of two batches of AAV-hSyn-Cre after injecting into the hypothalamus. It remains to be determined if the neurotoxicity stems from the AAV’s preparation in the particular batch. Moreover, it is essential to understand the impact of AAV-Cre on various glial cell types and neurons in different brain regions. Furthermore, the molecular mechanisms behind the neurotoxic effects have not been explored in this study, warranting future mechanistic research.

## Author contributions

R.S. conducted animal behavior assays, histology, and data analysis. E.P. helped with histology and imaging analysis. N.B. helped with histology and imaging analysis. Y.L. conducted stereotactic surgeries, animal behavior analysis, and histology. H.J.K. conducted CLAMS experiments and corresponding data analysis. L.W. conducted stereotactic surgeries. L.W. and Z.P.P. conceived the project, and designed the experiments, and wrote the paper with R.S. with input from all authors.

## Acknowledgments

We thank Dr. Larry Zweifel of the University of Washington for providing us the CRH^flox/flox^ mouse line. This study was supported by grants from the Robert Wood Johnson Foundation to the Child Health Institute of New Jersey (RWJF grant #74260), and the NIH NIDDK R01DK131452. L.W. was supported by the New Jersey Governor’s Council for Medical Research and Treatment of Autism Postdoctoral Fellowship (CAUT24DFP005) and the NExT-Metabolism Pilot Award (500301). R.S. was supported by Rutgers Aresty Research Center’s independent research award. The authors would like to thank Dr. Mark Rossi (Rutgers University) for critical reading of the manuscript and insightful comments. The graphical abstract was created with BioRender.

## Data Availability

The code used to analyze open-field test data can be found at https://github.com/RohanSavani/OpenFieldAnalysis. The data from the current study are available from the corresponding authors upon reasonable request.

## Supplementary Data

Supplementary data to this article: Supplemental Figures S1-S4

## Supplementary Information

**Figure S1:**
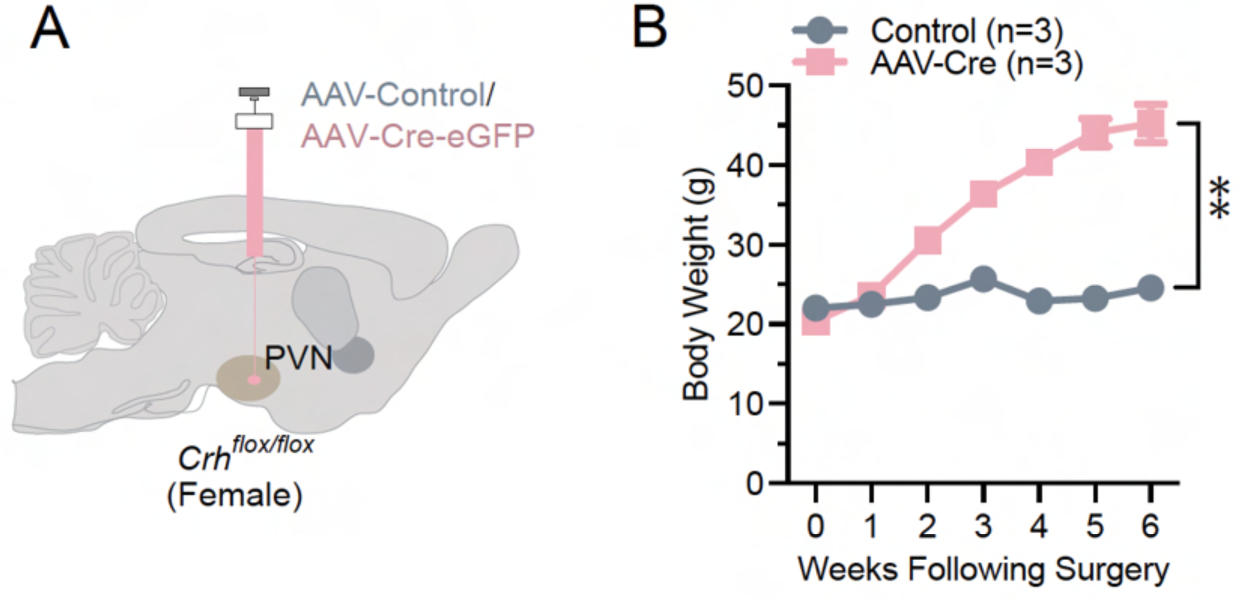
AAV-Cre expression in the PVN of female *Crh^f/f^* mice recapitulates body weight increases. **(A)** Experimental paradigm for virus delivery. **(B)** Body weight increases following AAV-Cre injection (two-way ANOVA, main effect of Group: F (1, 4) = 59.08, p = 0.0015; main effect of Time: F (1.782, 7.129) = 77.74, p < 0.0001; interaction between Group and Time: F (6, 24) = 58.41, p < 0.0001). Sample sizes are as shown on graphs (n = 3 mice per group). Data are presented as mean ± SEM.

**Figure S2:**
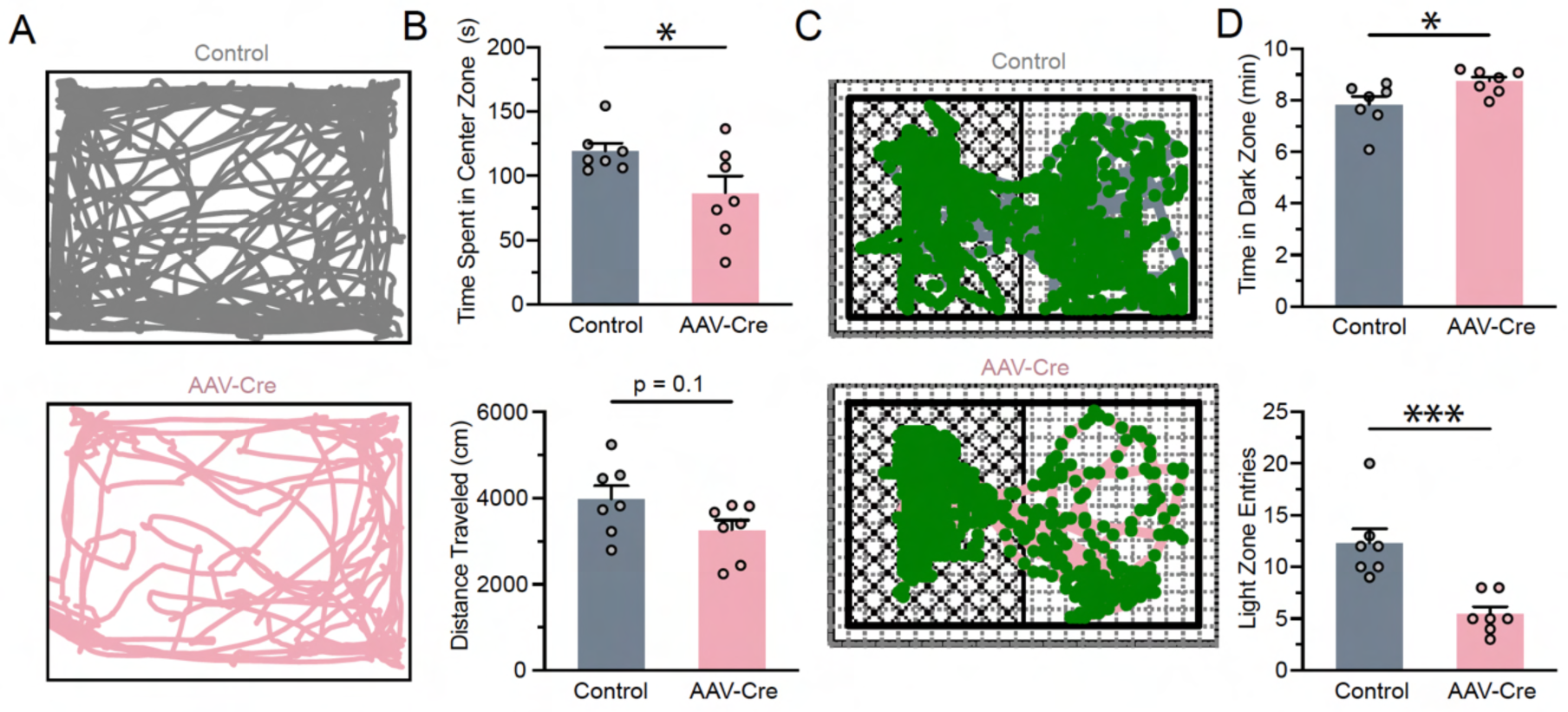
*Crh^f/f^* mice expressing AAV-Cre in PVN display elevated anxiety-like behaviors. **(A)** Representative trajectories of AAV-GFP-(top) and AAV-Cre-injected (bottom) male mice during the open field test. **(B)** AAV-Cre-injected mice spent less time in the center zone of the open field test (t (12) = 2.181, p = 0.0498), and changes in distance traveled were not significant (t (12) = 1.811, p = 0.0952) during the open field test. **(C)** Representative trajectories of both groups during the light-dark box test. **(D)** Time spent in the dark zone increased (t (12) = 2.416, p = 0.0326), and entries into the light zone decreased (Mann-Whitney test, U = 0, p = 0.0006) during the light-dark box test. Sample sizes are as shown on graphs (n = 7 mice per group). Data are presented as individual points and mean + SEM.

**Figure S3.**
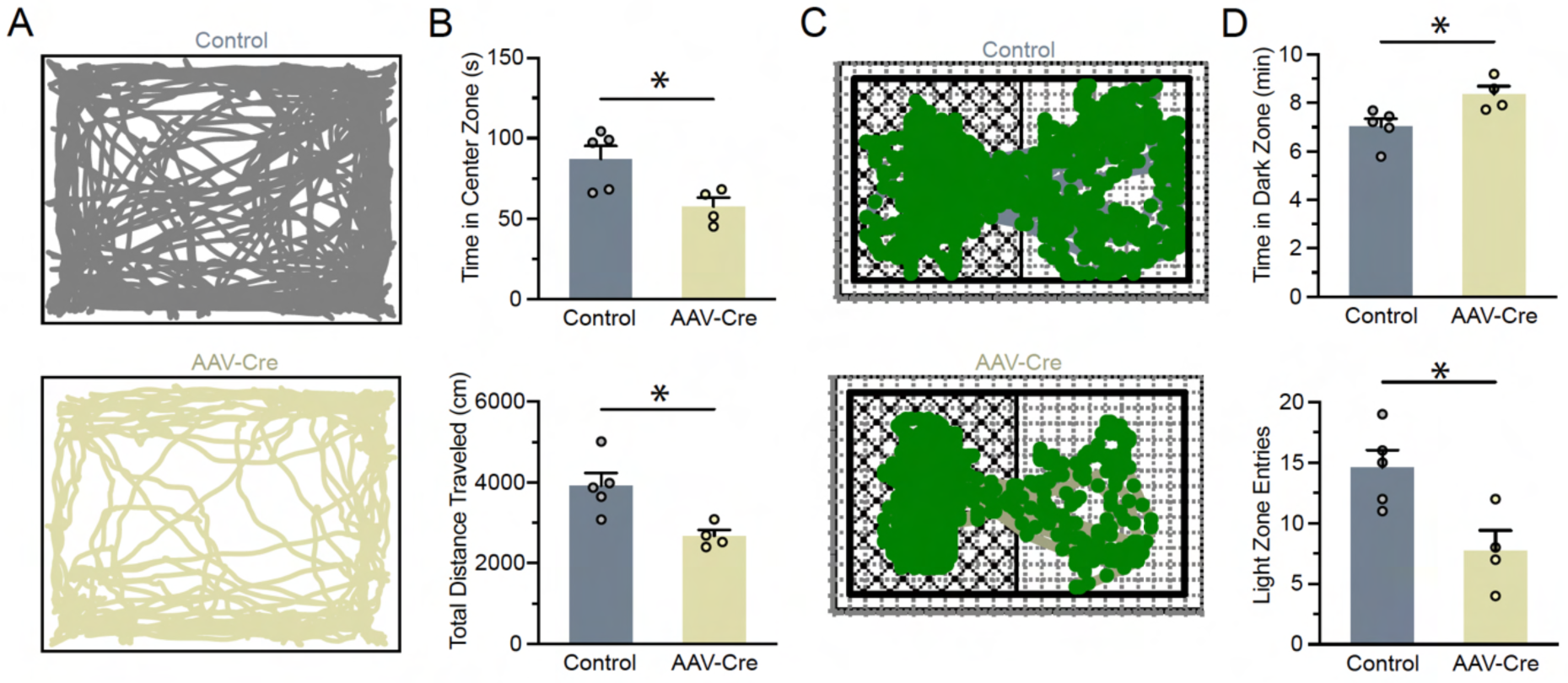
Wild-type male mice injected with AAV-Cre in PVN exhibit increased anxiety-like behaviors. **(A)** Representative traces of wild-type mice during the open field test. **(B)** During the open field test, time spent in the center zone decreased (t (7) = 2.817, p = 0.0259), and total distance traveled decreased (t (7) = 3.275, p = 0.0136). **(C)** Representative movement during the light-dark box test for both groups. **(D)** During the light-dark box test, time spent in the dark zone increased (t (7) = 2.798, p = 0.0266), and entries into the light zone decreased (t (7) = 3.142, p = 0.0163). Sample sizes are as shown on graphs (n = 5 AAV-GFP-injected mice, n = 4 AAV-Cre-injected mice). Data are presented as individual points and mean + SEM.

**Figure S4.**
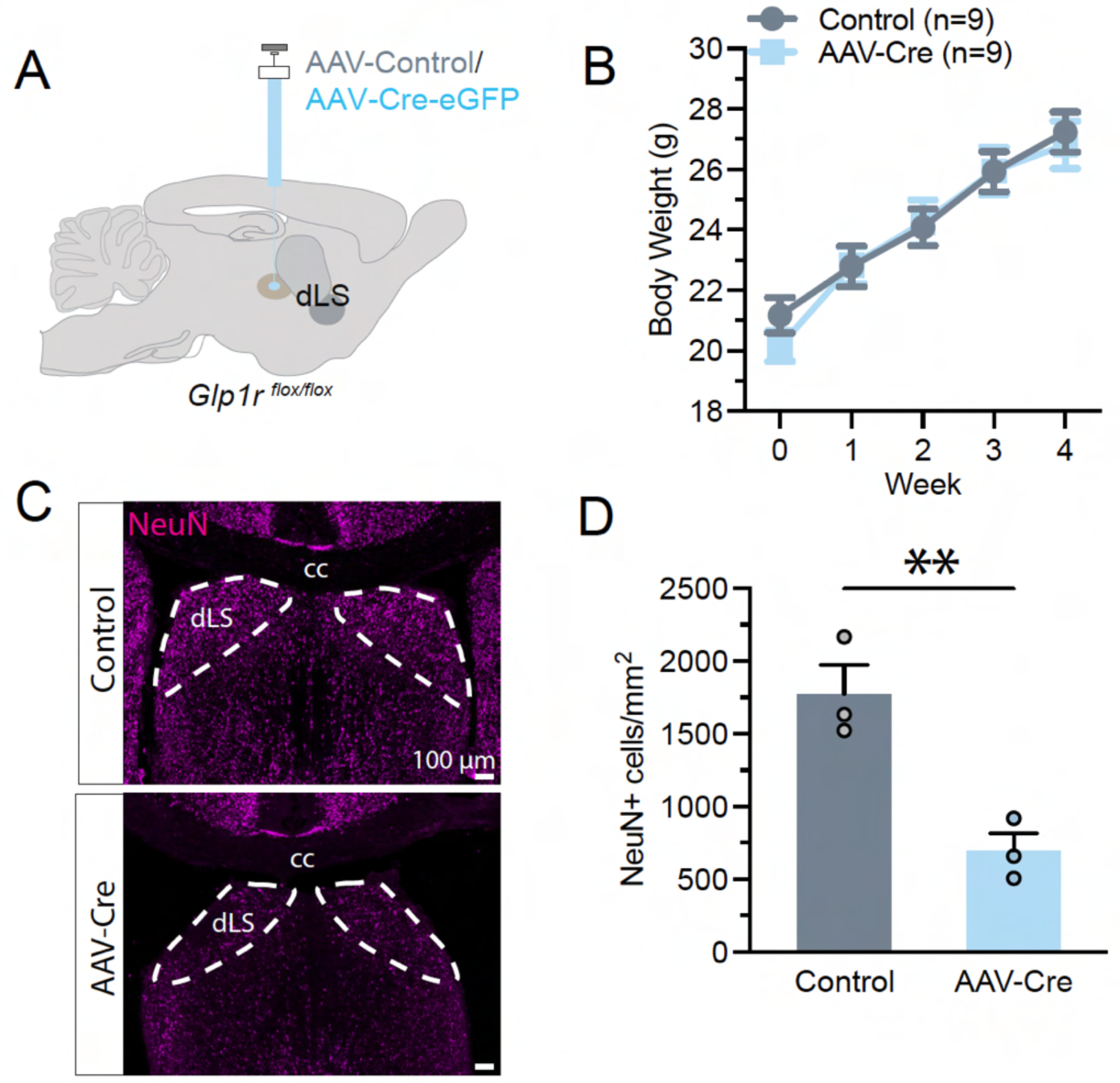
AAV-Cre expression in the dLS of male *Glp1r^f/f^* mice results in cell death without metabolic deficits. **(A)** Experimental paradigm for virus delivery. **(B)** Body weight was unchanged (two-way ANOVA, main effect of Group: F (1, 16) = 0.07282, p = 0.7907; main effect of Time: F (1.826, 29.22) = 364.6, p < 0.0001; interaction between Group and Time: F (4, 64) = 3.406, p = 0.0138). **(C)** Representative images of neuronal nuclei in the dLS after AAV-GFP or AAV-Cre injection. **(D)** Neuron density decreased in the dLS after AAV-Cre expression (t (4) = 4.631, p = 0.0098). Sample sizes are as shown on graphs: **(B)** n = 9 mice per group; **(D)** n = 3 mice per group. Data are presented as individual points and mean ± or + SEM. cc: corpus callosum.

**Figure S5:**
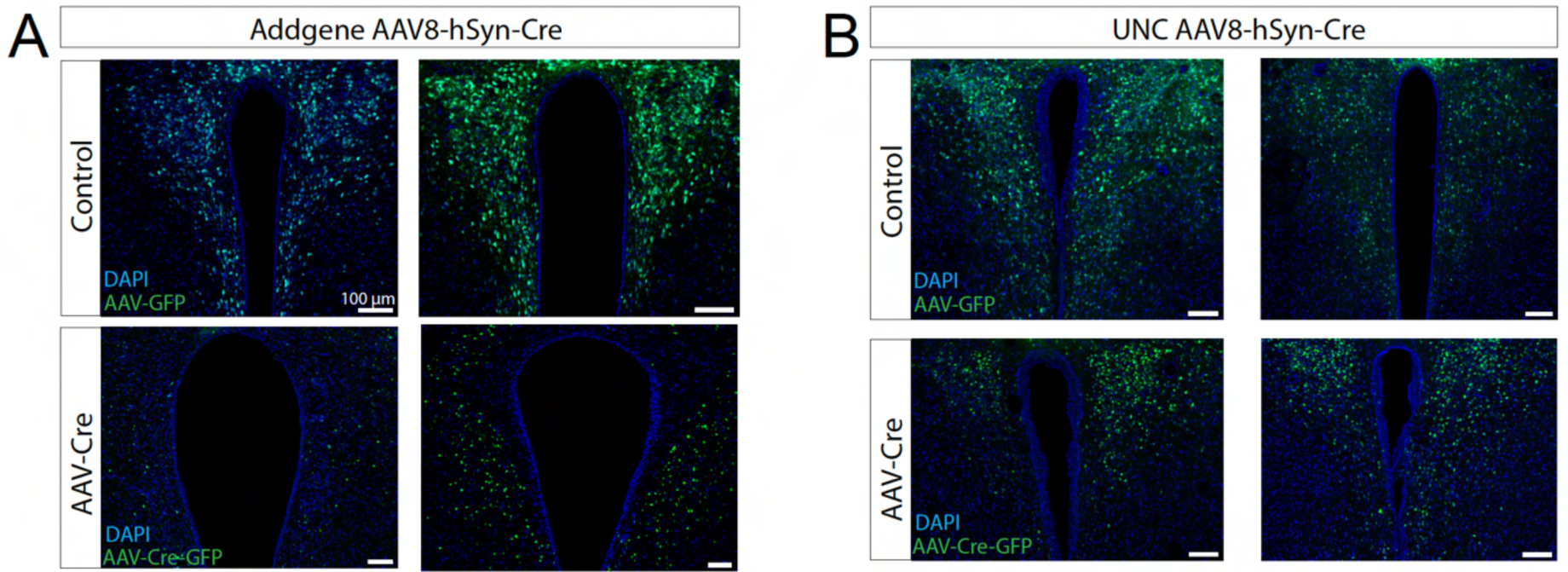
AAV-Cre-GFP Expression with AAV-Cre and AAV-GFP in PVN. **(A)** Examples of expression of AAV-GFP (control) or AAV-Cre-GFP in *Crh-flox* mice correspond to Figure 2. **(B)** Examples of expression in *Crh-flox* mice injected with the UNC Vector Core-sourced AAV-Cre correspond to Figure 4.

